# High-density lipoprotein characteristics and coronary heart disease: a Mendelian randomization study

**DOI:** 10.1101/673939

**Authors:** Albert Prats-Uribe, Sergi Sayols-Baixeras, Alba Fernández-Sanlés, Isaac Subirana, Robert Carreras-Torres, Gemma Vilahur, Fernando Civeira, Jaume Marrugat, Montserrat Fitó, Álvaro Hernáez, Roberto Elosua

## Abstract

**Background:** The causal role of high-density lipoproteins (HDL) in coronary artery disease (CAD) has been questioned. Our aim was to analyze whether genetically determined quantitative and qualitative HDL characteristics were independently associated with CAD.

**Methods:** We designed a two-sample multivariate Mendelian randomization study with available genome-wide association summary data. We identified genetic variants associated with HDL cholesterol and apolipoprotein A-I quantity, HDL size, particle levels, and lipid content to define our genetic instrumental variables in one sample (Kettunen et al study, *N*=24,925) and analyzed their association with CAD risk in a different study (CARDIoGRAMplusC4D, *N*=184,305). We validated these results by defining our genetic variables in another database (METSIM, *N*=8,372) and studied their relationship with CAD in the CARDIoGRAMplusC4D dataset. To estimate the effect size of the associations of interest adjusted for other lipoprotein traits (potential pleiotropy) we used the Multi-trait-based Conditional & Joint analysis.

**Results:** Genetically determined HDL cholesterol and apolipoprotein A-I levels were not associated with CAD. HDL mean diameter (β=0.27 [95%CI=0.19; 0.35]), cholesterol levels in very large HDLs (β=0.29 [95%CI=0.17; 0.40]), and triglyceride content in very large HDLs (β=0.14 [95%CI=0.040; 0.25]) were directly associated with CAD risk, whereas the cholesterol content in medium-sized HDLs (β=-0.076 [95%CI=-0.10; −0.052]) was inversely related to this risk. These results were validated in the METSIM-CARDIoGRAMplusC4D data. Genetic variants linked to both HDL qualitative traits and CAD risk were located within *LIPC*, the *APOE/C1/C4/C2* cluster, *APOB, PCIF1-PLTP*, and *TTC39B*.

**Conclusions:** Some HDL characteristics related to size, particle distribution, and triglyceride content are related to CAD risk whilst HDL cholesterol levels are not. This relationship could be mediated by the hepatic lipase; the apolipoproteins E, C-I, and B; the phospholipid transfer protein; and the tetratricopeptide repeat domain protein 39B, which arise as potential therapeutic targets in cardiovascular disease.

## INTRODUCTION

The inverse association between high-density lipoprotein cholesterol (HDL-C) levels and the risk of coronary artery disease (CAD) has been reported in observational studies^1^. However, experimental and genetic studies question the causality of this association. On the one hand, drugs such as fibrates, niacin, and cholesteryl ester transfer protein inhibitors increase HDL-C levels but do not decrease CAD risk^2^. On the other hand, genetic predisposition to high HDL-C levels has not been linked to any decrease in the risk of cardiovascular events^3,4^. Thus, researchers are looking beyond HDL-C levels to disentangle this apparent contradiction. Anti-atherogenic properties of HDL particles seem to be determined by the quality or function of the lipoprotein^5^. HDL particle size and number have been linked to cardiovascular risk^6^, and this association could be mediated through HDL functionality, which is predictive of cardiovascular risk^7^. The interplay between HDL-C and triglyceride levels, as two of the faces of atherogenic dyslipidemia, may also play a relevant role in their relationship with CAD^8,9^. Thus, further evidence of causal association between HDL characteristics and CAD risk would provide relevant data on the validity of these particles as therapeutic targets. Mendelian Randomization (MR) studies have arisen as a powerful tool to ascertain the potential causality of the association between a biomarker and a disease^10^. These studies assess the association between the genetically determined lifelong values of a biomarker and the development of a clinical outcome. MR studies have already raised serious doubts on the causal role of quantitative HDL characteristics, such as HDL-C and apolipoprotein A-I (ApoA-I) levels, in CAD^3,11^. However, to date, the association between qualitative HDL characteristics and CAD has not been tested using a MR approach. HDL mean diameter, the concentration of HDL particles of each size subtype, the distribution of cholesterol across the HDL size subtypes, and the presence of other lipids in HDL particles (such as triglycerides, highly present in large HDLs) are some of these qualitative traits. Additionally, this evaluation must take into account the complexity of lipid metabolism and its potential genetic pleiotropic effects. HDL-C, low-density lipoprotein cholesterol (LDL-C), and triglyceride levels are highly interdependent and, therefore, the method used to test the association between HDL properties and CAD risk should take into account this inter-correlation^12^. This study had two aims: 1) to assess the potential causal association of quantitative and qualitative HDL characteristics with CAD risk, using a two-sample MR approach; and 2) to explore potential mechanisms explaining the observed associations.

## METHODS

### Study design and data sources

We designed a two-sample MR study using aggregated summary data^10^ from three published meta-analyses of genome-wide association studies. The main analysis was based on data from Kettunen et al^13^ (*N*=24,925) and the CARDIoGRAMplusC4D consortium^14^ (*N*=184,305), and the validation analysis used the METSIM^15^ (*N*=8,372) and the CARDIoGRAMplusC4D datasets. A more detailed description of the studies is available in *Supplemental Materials*.

We centered our analysis on the genetic variants associated with: 1) the main lipid profile traits in serum (HDL-C, LDL-C, and triglyceride levels); 2) other measurements of HDL quantity (ApoA-I levels); 3) HDL mean diameter; 4) the quantities of cholesterol transported in small, medium-sized, large and very large HDLs; 5) the quantity of other lipid species in HDL particles (triglycerides transported in very large HDLs); and 6) the levels of HDL particles according to the previous HDL size subtypes. Both the Kettunen et al and the METSIM studies measured HDL qualitative characteristics by the same nuclear magnetic resonance spectroscopy technique^16^.

### Assessment of genetic variants linked to lipoprotein characteristics and CAD risk adjusted for other lipoprotein traits: Multi-Trait-based Conditional & Joint analyses

To identify the genetic variants associated with each lipoprotein characteristic that were also linked to CAD considering the potential pleiotropy among lipid profile traits, we used the Multi-trait-based Conditional & Joint analysis^17^ in both the main and the validation stage. This method enables the estimation of the magnitude of the association of each genetic variant with each lipoprotein characteristic and with CAD, independently from the other genetically determined lipoprotein traits (adjusted effect sizes). For example, if we considered HDL-C as “main variable” and LDL-C and triglyceride levels as “covariates”, we would obtain the betas and standard errors of the associations of genetic variants with (1) HDL-C (using the Kettunen and METSIM raw summary data) and (2) CAD (using the CARDIoGRAMplusC4D raw summary results), adjusted for the association of these same genetic variants with LDL-C and triglyceride concentrations. For this purpose, we defined six multivariate models a priori. Model 1 included HDL-C, LDL-C and triglyceride levels. Further models included the elements in Model 1 as covariates and the following parameters as main variables: ApoA-I levels (Model 2); HDL mean diameter (Model 3); the cholesterol content in each HDL size subtype (small, medium-sized, large, and very large HDL particles; Model 4); the levels of HDL particles of each size subtype (Model 5); and the triglyceride content in very large HDLs (Model 6). In model 4 and 5, we required for the presence of at least two of the HDL subtypes (small, medium-sized, large, and very large) traits to build the model. The genetic correlation between traits was estimated by linkage disequilibrium score regressions using all genetic variants.

### Mendelian randomization analyses

Based on the adjusted gene variant effects and their standard errors computed as previously described, we performed the MR analysis using the Generalized Summary-data-based Mendelian Randomization method^18^. The genetic variants to be considered were selected with the following criteria: 1) strong association with the lipid traits of interest (*p*-value<5·10^−8^); 2) not in linkage disequilibrium (R^2^<0.01, using the 1000 Genome project data –http://www.1000genomes.org/phase-3-structural-variant-dataset– as reference^19^); and 3) a minor allele frequency ≥0.05.

As an additional approach to exclude potential pleiotropy, we also removed the variants with a significant result in the HEIDI-outlier test (*p*-value<0.01). Finally, we explored and confirmed the exclusion of potential pleiotropic effects using Egger regressions^20^. Statistical significance of our results was corrected for multiple comparisons (*p*-value=0.05/number of traits). A description of complementary sensitivity analyses using other MR analyses methods such as the median-based and inverse variance weighted, using the Global Lipid Genetic Consortium dataset^21^ and *post-hoc* statistical power estimations^22^ is available in *Supplemental Materials*.

## RESULTS

### Mendelian randomization results

#### Selected genetic variants

We identified genetic variants significantly associated with 13 lipoprotein characteristics in the data published by Kettunen et al. and with 8 lipoprotein traits in the METSIM data. The number of genetic variants for each lipoprotein trait ranged from 6 (level of medium-sized HDL particles) to 22 (ApoA-I). Genetic variants included in the analyses and their unadjusted and adjusted effects are listed in **Supplemental Excel File 1**.

We observed high inverse genetic correlations (correlation coefficient≤-0.50) between triglyceride and HDL-C levels. Conversely, we observed very high direct genetic correlations (correlation coefficient≥0.70) between the cholesterol content in each HDL size subpopulation and HDL-C concentrations, between HDL mean diameter and the level of very large HDL particles, and between ApoA-I and the level of very large HDL particles and their cholesterol content (**Supplemental Figure 1**).

#### Main analysis

We defined as statistically significant those associations with a *p*-value<3.85·10^−3^ (0.05/13). We observed a direct association between CAD risk and genetically determined levels of LDL-C (β=0.26 [95% Confidence Interval]: 0.17; 0.35], *p*-value=1.32·10^−8^) and triglycerides (β=0.18 [0.073; 0.29], *p*-value=1.05·10^−3^). Conversely, the genetically determined concentrations of HDL-C (β=0.008 [−0.084; 0.099], *p*-value=0.871) or ApoA-I (β=0.060 [−0.015; 0.13], *p*-value=0.116) were not associated with CAD risk (**Figures 1, 2A and 2B**).

**Figure 1.**
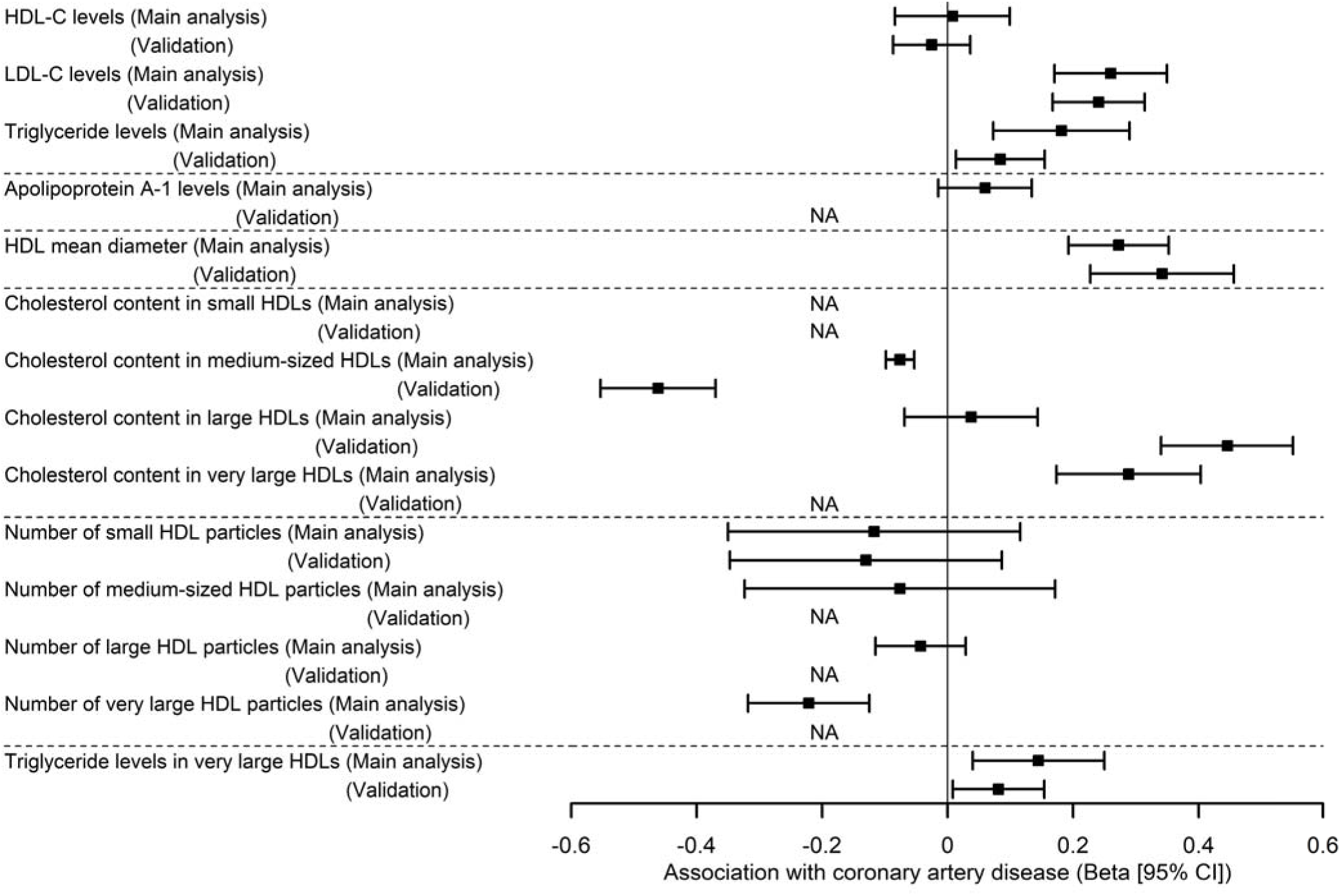
Association of the genetically determined lipid profile and HDL characteristics with coronary artery disease risk. Effect of genetically determined HDL cholesterol (HDL-C), LDL cholesterol (LDL-C) and triglyceride levels (Panel 1), apolipoprotein A-I concentrations (Panel 2), mean HDL diameter (Panel 3), cholesterol content in each HDL size subtype (Panel 4), number of particles of each HDL size subtype (Panel 5), and triglyceride levels in very large HDLs (Panel 6) on CAD, independently from the effect of genetically determined levels of classic lipid profile parameters. In all cases, main Mendelian randomization analyses (Kettunen-CARDIoGRAMplusC4D) appear first and the validation ones (METSIM-CARDIoGRAMplusC4D) appear below. NA: non-available association.

**Figure 2.**
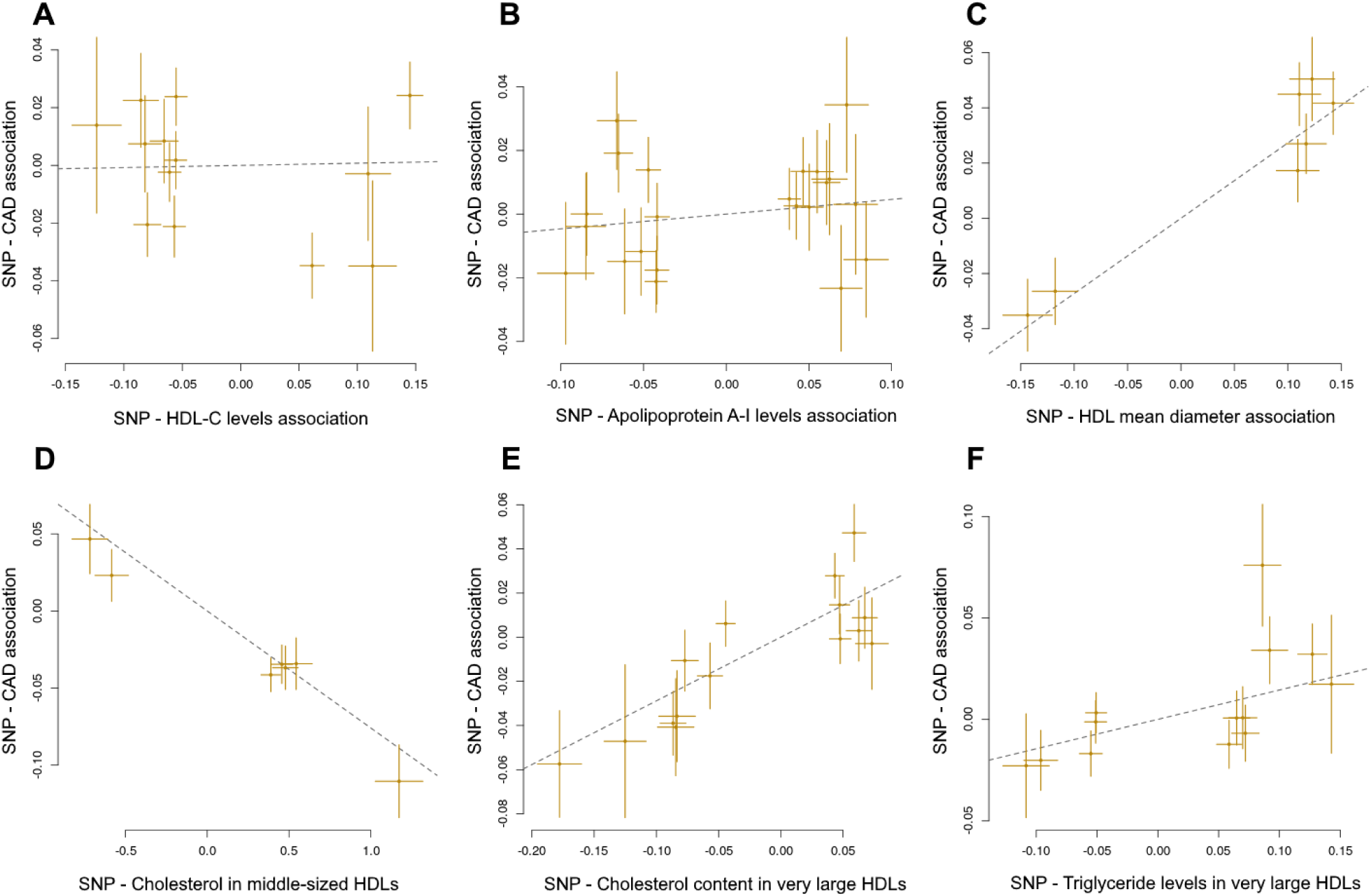
Association of individual SNPs affecting lipoprotein traits with coronary artery disease risk in the main analysis. Estimates of the associations of individual SNPs related to (A) HDL cholesterol (HDL-C) levels, (B) apolipoprotein A-I concentrations, (C) mean HDL diameter, (D) cholesterol content in medium-sized HDLs, (E) cholesterol content in very large HDLs, and (F) triglyceride levels in very large HDLs with coronary artery disease risk. Estimates were derived from the study by Kettunen et al and the CARDIoGRAMplusC4D meta-analyses (multivariate adjusted estimates). Error bars represent 95% confidence intervals. The slopes of the lines show the genetic instrumental variable regression estimates of the effect of the lipid characteristics on coronary artery disease risk.

In qualitative HDL measurements, the genetically determined HDL mean diameter was directly associated with CAD risk (β=0.27 [0.19; 0.35], *p*-value=2.23·10^−11^) (**Figures 1 and 2C**). Cholesterol levels in very large HDLs was also positively linked to CAD risk (β=0.29 [0.17; 0.40], *p*-value=8.90·10^−7^), whereas cholesterol in medium-sized HDLs was inversely related to this risk (β=-0.076 [−0.10; −0.052], *p*-value=4.55·10^−11^) (**Figures 1, 2D, and 2E**). The genetically determined levels of all subtypes of HDL particles showed an inverse trend towards an association with CAD risk, but only that between very large HDLs and CAD was statistically significant (β=-0.22 [−0.32; −0.12], *p*-value=7.12·10^−6^) (**Figure 1**). Finally, the genetically determined levels of triglycerides in very large HDLs were directly related to CAD risk (β=0.14 [0.040; 0.25], *p*-value=6.84·10^−3^) (**Figures 1 and 2F**). Effect sizes of all the associations are available in **Supplemental Table 1**.

Genetic variants linked to both HDL qualitative traits and CAD risk were located within *LIPC*, the *APOE/C1/C4/C2* cluster, *PCIF1, TTC39B*, and *APOB* (**Supplemental Excel File 1**).

#### Validation analysis

In the validation analysis (**Figure 1**), we confirmed the direct association between the genetically determined concentrations of LDL-C and CAD risk (β=0.24 [0.17; 0.31], *p*-value=1.21·10^−10^), triglycerides and CAD risk (β=0.084 [0.013; 0.15], *p*-value=0.020), and the null link between the genetically determined HDL-C levels and CAD (β=-0.025 [−0.087; 0.036], *p*-value=0.419).

Results regarding qualitative HDL traits were also replicated. We confirmed the direct association of genetically determined HDL mean diameter with CAD risk (β=0.34 [0.23; 0.46], *p*-value=4.47·10^−9^). There was a positive link between the cholesterol content in large HDLs and CAD (β=0.45 [0.34; 0.55], *p*-value=1.61·10^−16^) and an inverse relationship between the genetically determined cholesterol levels in medium-sized HDLs and CAD risk (β=-0.46 [−0.55; −0.37], *p*-value=5.9·10^−23^). Finally, high genetically determined levels of triglycerides in very large HDLs were nominally associated with greater CAD risk (β=0.081 [0.008; 0.15], *p*-value=0.030) (**Figure 1** and **Supplemental Figure 2**). We could not assess the associations between the levels of HDL particles and CAD due to the lack of genetic variants associated with at least two of these HDL traits in the METSIM study. The effect sizes of all the associations of interest are available in **Supplemental Table 2**.

#### Sensitivity analysis

Egger regression intercept estimates supported the absence of pleiotropic effects. Results of the median-based and inverse variance weighted methods confirmed the direction and significance of the main analyses (**Supplemental Table 3**).

Associations between genetically determined HDL-C, LDL-C and triglyceride levels and CAD risk identified in our main analysis were similar to those obtained from the Global Lipid Genetic Consortium (**Supplemental Table 4**).

#### Post-hoc statistical power estimation

Power estimation for the main analyses ranged from 2.4% to 96.9% (**Supplemental Table 5**).

## DISCUSSION

Our findings suggest a potential causal relationship between qualitative HDL characteristics and CAD risk, even though HDL-C and ApoA-I levels were not associated with CAD. In particular, genetically determined mean HDL size, the distribution of cholesterol across HDL size subpopulations, and the triglyceride content in HDL particles were related to CAD risk.

The relationship between HDL and cardiovascular risk is controversial^4^. Recent studies suggest that HDL functions and quality characteristics, rather than HDL-C concentration, are the main determinants of HDL anti-atherogenic properties^5^. Our data are consistent with previous evidence, and reflect that HDL-C and ApoA-I levels in the bloodstream are not causally related to CAD^3,11^. However, we observed a decrease in CAD risk when HDL-C was mainly transported in smaller HDLs, but an increase in CAD risk when HDL-C was carried by larger HDL particles (in both main and validation analyses, there is a gradient towards greater CAD risk as more cholesterol is transported in larger HDLs). Our results contribute to explaining why HDL-C levels are not causally associated with cardiovascular risk, as the protective effect of cholesterol content in medium-sized HDLs may be counterbalanced by the adverse effect of cholesterol levels in larger particles. These results are consistent with previous experimental evidence, and could contribute to explain the therapeutic failure of the pharmacological agents known to increase HDL-C levels. Niacin or cholesteryl ester transfer protein inhibitors are effective in increasing HDL-C concentrations but not in reducing CAD risk; this paradox could be explained by a promotion of the accumulation of cholesterol content in large HDLs^2^. In gemfibrozil-treated patients, changes in HDL-C levels accounted for a small proportion of the CAD risk reduction (<10%), whereas the increase in small HDLs was much more predictive of this risk reduction^23^. Our results also concur with genetic studies analyzing variants in the *SR-B1* gene, showing that individuals with loss-of-function variants have higher HDL-C concentrations, mainly in very large particles, but also higher CAD risk^24^.

However, there is still controversy in the relationship between HDL size subtypes and cardiovascular risk: some authors advocate for small HDLs as indicators of lower CAD risk^25^ while others suggest they are associated with increased CAD risk^26^. There are several possible explanations for this heterogeneity. First, baseline health conditions of the subjects affect HDL quality and function. Lipid-poor, protein-rich, small HDLs could be dysfunctional in pro-oxidative and pro-inflammatory pathological states due to post-translational modifications of their proteins and their enrichment in pro-inflammatory mediators (such as serum amyloid A or complement 3)^27^. Second, laboratory procedures to measure HDL size (nuclear magnetic resonance spectroscopy, electrophoresis, etc.) differed between the published studies, and there is low concordance between these techniques^28^. Third, the statistical models used did not consider all the same confounding factors and did not always include as covariates the levels of HDL-C or other lipid profile parameters related to these lipoproteins (e.g. triglyceride concentrations).

Triglyceride and HDL-C levels may be two sides of the same coin, and this relationship may contribute to explaining why HDL-C is not causally related to CAD while triglycerides are. Hypertriglyceridemic states (generally due to high levels of very-low density lipoprotein concentrations in plasma) are linked to an increased activity of the cholesteryl ester transfer protein, an enzyme that exchanges triglycerides in very-low density lipoproteins for cholesterol in HDLs, resulting in an enrichment of HDLs in triglycerides^8^. Aged HDL particles may also become increasingly richer in triglycerides because this exchange is an essential process by which HDLs get rid of the cholesterol they have collected from peripheral cells and transfer it back to the liver^29^. In any case, triglyceride-rich HDLs have been shown to present their ApoA-I in an unstable conformation^30^, which may be related to lower HDL function (lower cholesterol efflux capacity) and a greater disintegration of the HDL structure (lower HDL-C levels)^8^. Our results confirm that triglyceride-rich HDLs are causally related to higher CAD risk independently from the circulating levels of triglycerides and HDL-C. In addition, this mechanism also verifies the hypothesis that high triglycerides (in circulation and in HDL particles) are essential mediators of high cardiovascular risk and suggests that low HDL-C levels in these states may be a secondary consequence of this lipid disruption. Both observational and experimental studies have more consistently found an inverse relationship between the number of HDL particles and cardiovascular risk, compared to HDL-C levels^7^. Similarly, we observed that the concentrations of HDL particles of all sizes were inversely related to CAD risk, although only the genetically determined levels of very large HDLs were significantly associated with it in the main analysis. Unfortunately, we could not validate these results due to the lack of valid genetic variants in the METSIM study.

Our results are mechanistically plausible and highlight novel potential therapeutic targets in cardiovascular disease, since genetic variants individually associated with HDL qualitative traits and CAD in our data were located within five HDL-related genes or gene clusters. First, *LIPC* encodes for hepatic lipase C, an enzyme that hydrolyzes triglycerides in circulating lipoproteins, including HDL particles^29^. Hydrolysis of HDL triglycerides by this enzyme generates small/medium-sized, triglyceride-depleted particles, considered to be more stable and functional than very large, triglyceride-rich HDLs^31^. Since triglyceride-rich HDLs were also causally linked to CAD in our data, this potential mechanism would contribute to explaining a decrease in cardiovascular risk. Second, the *APOE/C1/C4/C2* cluster encodes apolipoproteins E, C-I, C-II, and C-IV and has been classically associated with blood lipid levels^32^. Among these proteins, two are particularly related to HDL metabolism: apolipoprotein E (ApoE) and apolipoprotein C-I (ApoC-I). ApoE is a pivotal mediator in reverse cholesterol transport which is one of the most essential HDL function^33^. ApoC-I is involved in the modulation of two key enzymes of HDL cholesterol metabolism: the activation of lecithin-cholesterol acyltransferase and the inhibition of cholesteryl ester transfer protein^34,35^. The third HDL-related locus encodes the PDX1 C-Terminal Inhibiting Factor 1 and is located next to the *PLTP* gene, which expresses the phospholipid transfer protein, an enzyme involved in HDL remodeling/stabilization and the generation of lipid-free/lipid-poor small HDLs^29^. *PCIF1*-related gene variants have been shown to modulate phospholipid transfer protein function in other studies^36^. Fourth, *TTC39B* encodes the tetratricopeptide repeat domain protein 39B, whose genetic variants had already been associated with HDL-C levels and CAD in previous works^32,37^. Fifth, *APOB* encodes for apolipoprotein B, the essential apolipoprotein in chylomicrons, LDLs, and very-low density lipoproteins. Although its relationship with HDL biology may seem distant, gene variants located in this gene have been surprisingly associated with HDL cholesterol levels in four large genome-wide association studies^32,38–40^.

Our study has several methodological strengths. First, our results are based in MR, a useful approach to explore the causality of the association between biomarkers and specific diseases. Second, we included two independent MR analyses to validate the results initially observed. Finally, the validity of the genetic variants for HDL-C, LDL-C and triglyceride levels initially generated was confirmed, supporting the validity of these datasets for the analysis of other genetic variants. However, our study also has limitations. First, in order to use a MR approach, we had to make some assumptions^10^, among which stands out the absence of pleiotropy. In our case, most of the genetic variants used as instruments were associated with more than one lipid trait. To solve this problem, we used a novel approach (Multi-trait based Conditional & Joint analysis– Generalized Summary data-based Mendelian randomization methodology) to control for the confounding effects related to the close relationship between lipoprotein characteristics and to minimize pleiotropy^17^. Second, the interpretation of multivariable MR is challenging, especially when the covariate-biomarker lies on the causal pathway from the main-biomarker to disease, or when the covariate-biomarker measures the same entity as the main-biomarker^41^. Third, the statistical power of our analyses was limited for some of the traits of interest. Finally, in the validation analysis we could not generate GIVs for some of the lipoprotein traits.

## CONCLUSIONS

Our study indicates that several genetically and life-long qualitative HDL characteristics were related to CAD risk. Although HDL-C and ApoA-I levels were not causally linked to CAD, our results support a potential causality between higher mean HDL diameter, greater cholesterol levels in very large HDLs, and triglyceride-rich HDL particles and higher CAD risk, and between cholesterol levels in medium-sized HDLs and lower CAD risk. This relationship could be mediated by several HDL-related proteins (hepatic lipase; apolipoproteins E, C-I, and B; phospholipid transfer protein; and tetratricopeptide repeat domain protein 39B), which are suggested as potential therapeutic targets for further exploration in cardiovascular prevention.

## ACKNOWLEDGEMENTS

We thank Elaine M. Lilly, PhD, for her critical reading and revision of the English text. Data were downloaded from: www.computationalmedicine.fi/data#NMR_GWAS (Kettunen study), http://csg.sph.umich.edu/boehnke/public/metsim-2017-lipoproteins/ (METSIM study), www.cardiogramplusc4d.org (CARDIoGRAMplusC4D), http://csg.sph.umich.edu/abecasis/public/lipids2013/ (Global Lipid Genetic Consortium), http://www.1000genomes.org/phase-3-structural-variant-dataset (1000 Genome).

## SOURCES OF FUNDIND

This work was supported by the Instituto de Salud Carlos III–European Regional Development Fund (CD17/00122, IFI14/00007, PI18/00017), the Medical Research Council (MR/K501256/1, MR/N013468/1), the Spanish Ministry of Economy and Competitiveness (BES-2014–069718, SAF2015-71653-R), the European Union’s Horizon 2020 Research and Innovation Programme (796216), and the Government of Catalonia through the Agency for Management of University and Research Grants (2017 SGR 222). CIBERCV, CIBERESP and CIBEROBN are initiatives of Instituto de Salud Carlos III (Madrid, Spain).

