## Supplemental file for "High-density lipoprotein characteristics and coronary heart disease: a Mendelian randomization study"

**SUPPLEMENTAL MATERIAL**

### TABLE OF CONTENTS


1. **Supplemental Methods**
    - Genome-wide data on lipoprotein traits
    - Genome-wide data on coronary artery disease risk
    - Sensitivity analysis
    - Post-hoc statistical power estimation
2. **Supplementary Tables**
    - Supplemental Table 1: Results of the main Mendelian randomization meta-analyses (Kettunen et al-CARDIoGRAMplusC4D).
    - Supplemental Table 2: Results of the validation Mendelian randomization meta-analyses (METSIM-CARDIoGRAMplusC4D).
    - Supplemental Table 3: Results of the sensitivity analyses of the Mendelian randomization using the inverse variance weighted method, the weighted median method, and Egger regression for the main lipoprotein traits identified to be potentially causally related to CAD in the main and the validation analyses.
    - Supplemental Table 4: Association of the genetic instruments related to HDL cholesterol, LDL cholesterol, and triglycerides levels (identified in the Global Lipids Genetic Consortium) with CAD risk in the CARDIoGRAMplusC4D consortium
    - Supplemental Table 5: Post-hoc power estimation (for an odds ratio of 0.9, 1.1) for the genetic instrumental variables of the lipoprotein traits of interest.
3. **Supplementary Figures**
    - Supplemental Figure 1: Heat map of correlations between the genetic instrumental variables obtained in the Kettunen et al study.

- Supplemental Figure 2: Association of individual SNPs related to HDL traits with CAD risk in the validation analyses (METSIM-CARDIoGRAMplusC4D).

**4. Supplemental Excel Files**

- Supplemental Excel File 1: Genetic variants included in each genetic instrumental variable. Multi-trait based Conditional & Joint analysis output (multivariate adjusted association) and original data (univariate association).

### SUPPLEMENTAL METHODS

#### Genome-wide data on lipoprotein traits

The meta-analyses performed by Kettunen et al[13] included 14 European cohorts comprising 24,925 individuals. They tested the associations between 39 million genetic variants and 123 blood metabolites, 81 of them being lipid traits. Lipoprotein characteristics were determined from fasting serum samples using a standardized and previously described quantitative high-throughput nuclear magnetic resonance metabolomics methodology[16]. A regression method was used to estimate fasting state values in non-fasting samples. Data were downloaded from: www.computationalmedicine.fi/data#NMR_GWAS.

The METSIM study[15] involved 8,372 Finnish men and analyzed the associations between 15.1 million genetic variants and 71 lipid traits measured through proton nuclear magnetic resonance metabolomics using an additive genetic model. In this case, data were downloaded from: http://csg.sph.umich.edu/boehnke/public/metsim-2017-lipoproteins/.

Original studies complied with the Declaration of Helsinki, local ethics committee approved their respective research protocols, and their participants provided informed consent before joining the studies.

#### Genome-wide data on coronary artery disease risk

The CARDIoGRAMplusC4D study comprised 60,801 coronary artery disease (CAD) cases and 123,504 controls from 48 studies and analyzed 8.6 million genetic variants[14]. CAD was defined by the presence of myocardial infarction, acute coronary syndrome, chronic stable angina, or coronary stenosis >50%, and this definition could vary among the 48 studies included in the meta-analysis. Study data were downloaded from: http://www.cardiogramplusc4d.org0.

Original studies complied with the Declaration of Helsinki, local ethics committee approved their respective research protocols, and their participants provided informed consent before joining the studies.

**Sensitivity analysis**

We used inverse variance weighted methods, median based, and Egger regression to assess the consistency of the results, using the multivariate adjusted regression coefficients obtained with the Multi-trait based Conditional & Joint analysis. We also assessed whether the effect size of the association between the genetic variants of the main lipid traits and CAD was consistent using different datasets. We used the same methodology to identify the genetic variants for HDL-C, LDL-C and triglyceride levels in the dataset from the Global Lipid Genetic Consortium, which includes 188,577 individuals (http://csg.sph.umich.edu/abecasis/public/lipids2013/)[21]. Finally, we analyzed the association between those genetic variants and CAD risk in the CARDIoGRAMplusC4D Consortium data.

**Post-hoc statistical power estimation**

We calculated the power of our analyses to estimate associations between genetic variants and CAD risk using the method proposed by Brion et al[22]. We assumed that the effect size of the association was of a ±10% change in CAD risk (odds ratio of CAD 0.9 or 1.1). We also considered an alpha risk corrected for multiple comparisons. We estimated the variance of the lipoprotein traits explained by our GIVs as a sum of the genetic variances explained by each genetic variant selected.

**SUPPLEMENTARY TABLES**

**Supplemental Table 1.** Results of the main Mendelian randomization meta-analyses (Kettunen et al + CARDIoGRAMplusC4D).

| Trait | Number of SNPs | Beta coefficient | Standard error | *p*-value |
|---|---|---|---|---|
| HDL cholesterol levels | 13 | 0.008 | 0.047 | 0.87 |
| LDL cholesterol levels | 19 | 0.26 | 0.046 | $1.3 \cdot 10^{-8}$ |
| Triglyceride levels | 11 | 0.18 | 0.055 | $1.1 \cdot 10^{-3}$ |
| Apolipoprotein A-I levels | 22 | 0.046 | 0.049 | 0.34 |
| Mean HDL diameter | 7 | 0.27 | 0.041 | $2.2 \cdot 10^{-11}$ |
| Cholesterol content in small HDLs | - | - | - | - |
| Cholesterol content in medium-sized HDLs | 7 | -0.076 | 0.012 | $4.6 \cdot 10^{-11}$ |
| Cholesterol content in large HDLs | 7 | 0.038 | 0.054 | 0.49 |
| Cholesterol content in very large HDLs | 15 | 0.29 | 0.059 | $8.9 \cdot 10^{-7}$ |
| Number of small HDL particles | 8 | -0.12 | 0.12 | 0.32 |
| Number of medium-sized HDL particles | 6 | -0.076 | 0.13 | 0.55 |
| Number of large HDL particles | 7 | -0.043 | 0.037 | 0.24 |
| Number of very large HDL particles | 9 | -0.22 | 0.049 | $7.1 \cdot 10^{-6}$ |
| Triglyceride levels in very large HDLs | 13 | 0.14 | 0.054 | $6.8 \cdot 10^{-3}$ |

Associations of genetically determined HDL cholesterol, LDL cholesterol, and triglyceride levels with CAD are independent from the effect of the genetically determined levels of the rest of lipid parameters. Associations of genetically determined HDL qualitative traits are independent from the effect of the genetically determined levels of HDL cholesterol, LDL cholesterol, and triglycerides.

**Supplemental Table 2.** Results of the validation Mendelian randomization meta-analyses (METSIM + CARDIoGRAMplusC4D).

| Trait | Number of SNPs | Beta coefficient | Standard error | *p*-value |
|---|---|---|---|---|
| HDL cholesterol levels | 13 | -0.025 | 0.031 | 0.41 |
| LDL cholesterol levels | 13 | 0.24 | 0.037 | $1.2 \cdot 10^{-10}$ |
| Triglyceride levels | 9 | 0.084 | 0.036 | 0.02 |
| Apolipoprotein A-I levels | - | - | - | - |
| Mean HDL diameter | 3 | 0.34 | 0.058 | $4.5 \cdot 10^{-9}$ |
| Cholesterol content in small HDLs | - | - | - | - |
| Cholesterol content in medium-sized HDLs | 5 | -0.46 | 0.047 | $5.9 \cdot 10^{-23}$ |
| Cholesterol content in large HDLs | 7 | 0.44 | 0.054 | $1.6 \cdot 10^{-16}$ |
| Cholesterol content in very large HDLs | - | - | - | - |
| Number of small HDL particles | 3 | -0.13 | 0.111 | 0.23 |
| Number of medium-sized HDL particles | - | - | - | - |
| Number of large HDL particles | - | - | - | - |
| Number of very large HDL particles | - | - | - | - |
| Triglyceride levels in very large HDLs | 15 | 0.08 | 0.037 | 0.03 |

Associations of genetically determined HDL cholesterol, LDL cholesterol, and triglyceride levels with CAD are independent from the effect of the genetically determined levels of the rest of lipid parameters. Associations of genetically determined HDL qualitative traits are independent from the effect of the genetically determined levels of HDL cholesterol, LDL cholesterol, and triglycerides.

**Supplemental Table 3.** Results of the sensitivity analyses of the Mendelian randomization using the inverse variance weighted method, the weighted median method, and Egger regression for the main lipoprotein traits identified to be potentially causally related to CAD in the main and the validation analyses.

| | | Egger | | Inverse variance weighted method | | Weighted median method | | Egger regression | |
|---|---|---|---|---|---|---|---|---|---|
| Trait | Number of SNPs | Intercept | P-value | Regression coefficient | P-value | Regression coefficient | P-value | Regression coefficient | P-value |
| Kettunen et al + CARDIoGRAMplusC4D | | | | | | | | | |
| HDL mean diameter | 7 | -0.005 | $9.2 \cdot 10^{-1}$ | 0.274 | $6.4 \cdot 10^{-14}$ | 0.242 | $9.4 \cdot 10^{-6}$ | 0.312 | $4.0 \cdot 10^{-1}$ |
| Cholesterol in Medium-sized HDL particles | 7 | 0.001 | $9.6 \cdot 10^{-1}$ | -0.078 | $3.7 \cdot 10^{-14}$ | -0.076 | $3.9 \cdot 10^{-8}$ | -0.079 | $5.2 \cdot 10^{-2}$ |
| Cholesterol in Very large HDL particles | 15 | -0.007 | $5.7 \cdot 10^{-1}$ | 0.291 | $5.6 \cdot 10^{-06}$ | 0.313 | $1.5 \cdot 10^{-4}$ | 0.389 | $5.2 \cdot 10^{-2}$ |

**Supplemental table 4.** Association of the genetic instruments related to HDL cholesterol, LDL cholesterol, and triglycerides levels (identified in the Global Lipids Genetic Consortium) with CAD risk in the CARDIoGRAMplusC4D consortium.

| Trait | Number of SNPs | Beta coefficient | Standard error | *p*-value |
|---|---|---|---|---|
| HDL cholesterol levels | 61 | 0.001 | 0.031 | 0.97 |
| LDL cholesterol levels | 56 | 0.392 | 0.020 | $4.8 \cdot 10^{-89}$ |
| Triglyceride levels | 47 | 0.216 | 0.043 | $4.6 \cdot 10^{-7}$ |

**Supplemental Table 5**. Post-hoc power estimation (for an odds ratio of 0.9, 1.1) for the genetic instrumental variables of the lipoprotein traits of interest.

| Trait | Variability explained | Power |
|---|---|---|
| HDL cholesterol levels | 2.8% | 63.8% |
| LDL cholesterol levels | 3.9% | 82.7% |
| Triglyceride levels | 2.3% | 53.3% |
| Apolipoprotein A-I levels | 3.6% | 78.4% |
| Mean HDL diameter | 6.0% | 96.9% |
| Cholesterol content in small HDLs | - | - |
| Cholesterol content in medium-sized HDLs | 0.2% | 2.4% |
| Cholesterol content in large HDLs | 2.9% | 65.9% |
| Cholesterol content in very large HDLs | 2.2% | 49.0% |
| Number of small HDL particles | 0.3% | 3.9% |
| Number of medium-sized HDL particles | 0.6% | 9.1% |
| Number of large HDL particles | 1.4% | 29.1% |
| Number of very large HDL particles | 2.1% | 47.4% |
| Triglyceride levels in very large HDLs | 2.1% | 40.8% |

**SUPPLEMENTAL FIGURES**

**Supplemental Figure 1.** Heat map of correlations between the genetic instrumental variables obtained in the Kettunen et al study.

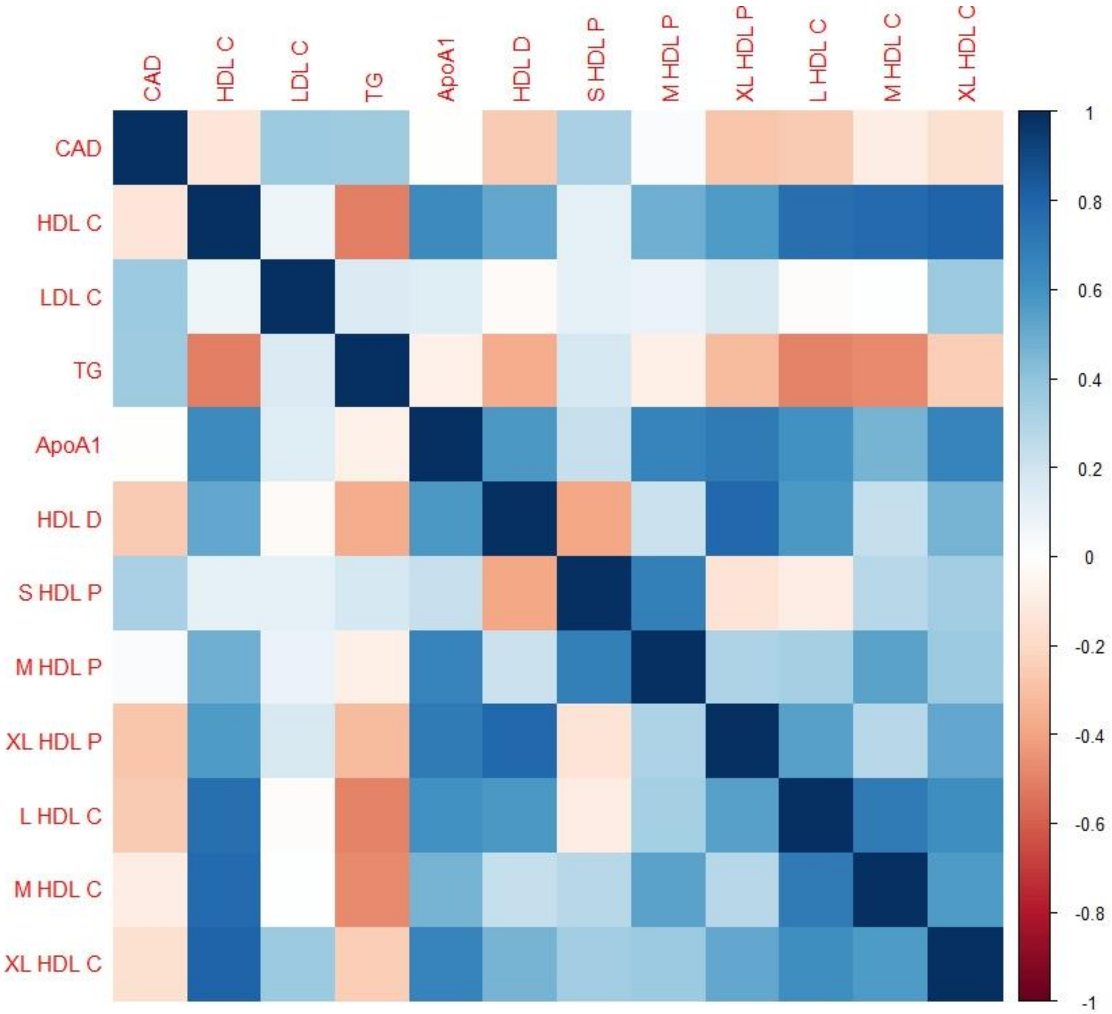


*CAD*: Coronary Artery Disease; *HDL-C*: high-density lipoprotein cholesterol; *LDL-C*: low-density lipoprotein cholesterol; *TG*: triglycerides; *ApoA-I*: apolipoprotein A-I; *HDL D*: mean HDL diameter; *S HDL P*: levels of small HDL particles; *M HDL P*: levels of medium-sized HDL particles; *XL HDL P*: levels of very large HDL particles; *L HDL C*: cholesterol content in large HDLs; *M HDL C*: cholesterol content in medium-sized HDLs; *XL HDL C*: cholesterol content in very large HDLs.

**Supplemental Figure 2.** Association of individual SNPs affecting lipoprotein traits with CAD risk in the validation analysis.

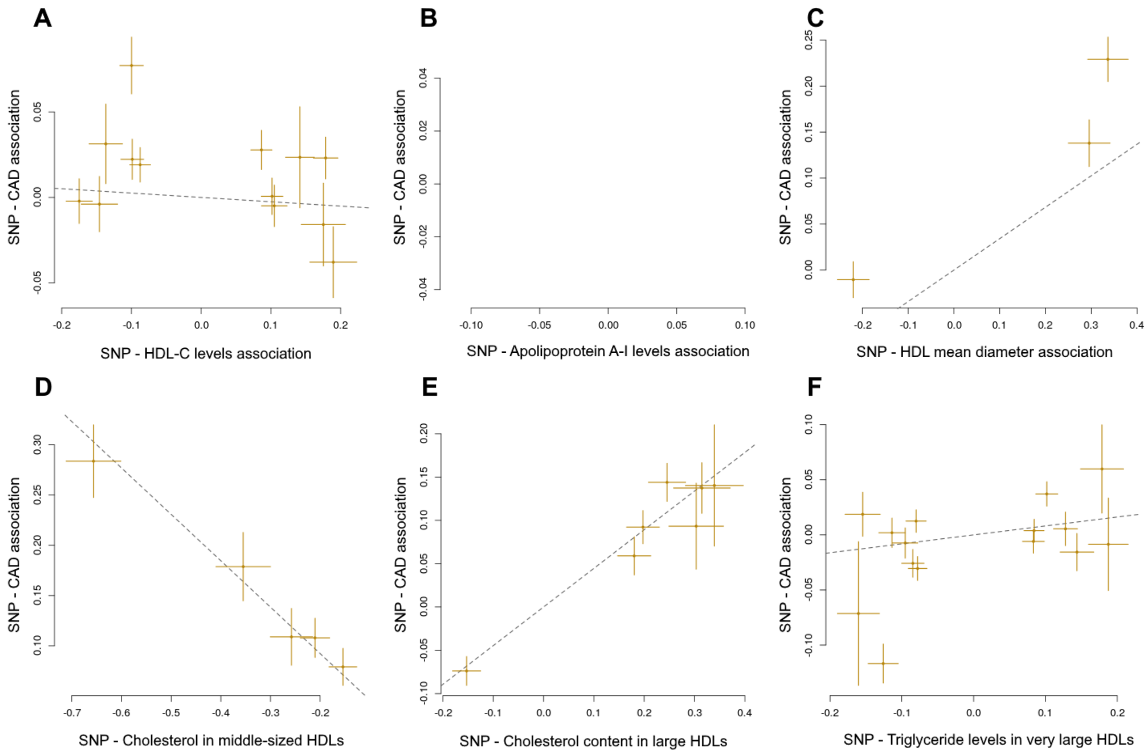


Estimates of the associations of individual SNPs related to (A) HDL cholesterol (HDL-C) levels, (B) apolipoprotein A-I concentrations, (C) mean HDL diameter, (D) cholesterol content in medium-sized HDLs, (E) cholesterol content in very large HDLs, and (F) triglyceride levels in very large HDLs with coronary artery disease risk. Multivariate adjusted estimates were derived from the METSIM-CARDIoGRAMplusC4D meta-analyses. Error bars represent 95% confidence intervals. The slopes of the lines show the genetic instrumental variable regression estimates of the effect of the lipid characteristics on coronary artery disease risk.

**Supplemental Excel File 1**. Genetic variants included in each genetic instrumental variable. Multi-trait based Conditional & Joint analysis output (multivariate adjusted association) and original data (univariate association).
